# hilldiv3: a unified Hill-number framework for neutral, phylogenetic and functional diversity analysis in R

**DOI:** 10.64898/2026.11.12.766119

**Authors:** Antton Alberdi, M Thomas P Gilbert, Ostaizka Aizpurua

## Abstract

1. Hill numbers provide a coherent family of diversity measures that can incorporate phylogenetic relatedness and functional trait dissimilarity, but applying them consistently across neutral, phylogenetic and functional diversity, and across measurement, partitioning and (dis)similarity, still requires combining several packages with incompatible conventions.
2. We present ‘hilldiv3’, an R package that provides per-sample diversity, nested multiplicative partitioning across scales for all three diversity types, bounded (dis)similarities and pairwise matrices, diversity profiles, evenness and redundancy, and the allocation of Hill power sums and dissimilarities to taxa or taxonomic sets.
3. ‘hilldiv3’ directly supported all twelve benchmarked combinations of operation and diversity type, and built pairwise (dis)similarity matrices 15–820 times faster than hilldiv2 and hillR, with up to two orders of magnitude less memory. We showcased the package by analysing Microflora Danica data (360 samples, 5,518 genomes, eight habitats), demonstrating that neutral, phylogenetic and functional diversity turned over at very different rates between habitats, and a near-constant Actinomycetota share concealed opposing shifts of two orders.
4. By deriving every metric from the same Hill-number formulation, ‘hilldiv3’ makes neutral, phylogenetic and functional results directly comparable, and lowers the barrier to rigorous diversity analysis of community-ecology and high-throughput sequencing data.

## Introduction

Quantifying the diversity of biological communities is a recurring task across ecology, evolution and, increasingly, microbiome science (Gotelli & Colwell, 2001). High-throughput sequencing now routinely produces feature tables of operational taxonomic units (OTUs), amplicon sequence variants (ASVs) or metagenome-assembled genomes (MAGs), which hold hundreds to thousands of features across many samples, and from which researchers compute a range of diversity metrics (Ji et al., 2013; Neu et al., 2021). Historically, the proliferation of such metrics has hampered comparability: classical indices are grounded in different mathematical bases, sit on different and non-linear scales, and respond differently to sampling effort, so values from one study often cannot be placed alongside those from another (Alberdi & Gilbert, 2019; Chao et al., 2014).

Hill numbers resolve much of this confusion by providing a single mathematical framework that encompasses many of these metrics as special cases (Hill, 1973; Jost, 2006). They express diversity as an effective number of units, that is, the number of equally abundant, equally distinct types (e.g., ASVs, MAGs, species) that would reproduce the observed diversity. A single order parameter, q, controls how strongly rare and common features are weighted. Reporting diversity across a range of q values therefore characterises a community without committing to a single, arbitrary index. In its neutral form it treats every feature as equally distinct. The phylogenetic form additionally weights diversity by evolutionary relatedness, so that a community of distant relatives counts as more diverse than one of close relatives (Chao et al., 2010); the functional form does the same using dissimilarity in measured traits, capturing differences in ecological role (Chao et al., 2014). Because all three rest on the same definition, their results are directly comparable. The framework also decomposes diversity multiplicatively into alpha (the average diversity within samples), gamma (the diversity of the pooled samples) and beta (their ratio, a measure of how much communities differ in composition) (Chao et al., 2016).

Alberdi & Gilbert reviewed the application of this framework to DNA-based data and argued for its routine use (Alberdi & Gilbert, 2019). In practice, however, assembling a complete Hill-number analysis still means combining several packages (e.g., hillR (Li, 2018), entropart (Marcon & Hérault, 2015), BAT (Cardoso et al., 2015) and vegan (Oksanen et al., 2013)), each with its own input conventions, alignment assumptions and output formats, and none of them supports all of these operations directly across the three diversity types. The limited computational efficiency of existing packages may result in excessively long analysis times, especially when performing phylogenetic and functional analyses on large datasets with many samples and taxonomic units.

To address these challenges, we developed ‘hilldiv3’ as a ground-up redesign of the ‘hilldiv’ lineage that computes a variety of neutral, phylogenetic and functional diversity analyses from a single Hill-number engine. Beyond consolidating existing functionality, ‘hilldiv3’ adds nested multiplicative partitioning that spans all three diversity types, and the attribution of Hill power sums and dissimilarities to individual taxa or taxonomic sets. Here, we describe the package’s structure and usage, benchmark it against existing implementations, and demonstrate it with a comprehensive diversity analysis of Danish environmental microbiomes from Microflora Danica (Singleton et al., 2026).

## Methods

### Design and architecture

‘hilldiv3’ separates the mathematics from the user interface. The numerical core, including per-sample alpha, the alpha/gamma/beta partition, and the beta-to-(dis)similarity conversions, lives in an internal compute engine that is unit-tested independently against hand-computed constants, and across edge cases (single taxon, empty sample, the q = 1 limit). The user-facing functions (hilldiv(), hillpart(), hilldiss(), hillsim(), hillpair(), hillprof(), hilleven(), hillred(), hillshare() and hillcontrib()) are thin wrappers around that engine. This separation makes the mathematic formulation auditable and keeps the public API stable.

All inputs pass through one validation and alignment layer: as_hill_input() accepts the various object types and match_data() reconciles their labels, comparing names as sets and reordering the count table to match the supplied tree or distance matrix. A mismatch is therefore corrected rather than silently computed on. This is a common failure mode of stitched-together workflows, including the previous version of this package. The same adapter accepts ordinary matrices, data frames and tibbles as well as phyloseq (McMurdie and Holmes 2013) and TreeSummarizedExperiment (Huang et al. 2020) objects, so the package slots into existing microbiome pipelines without reshaping.

**Figure 1.**
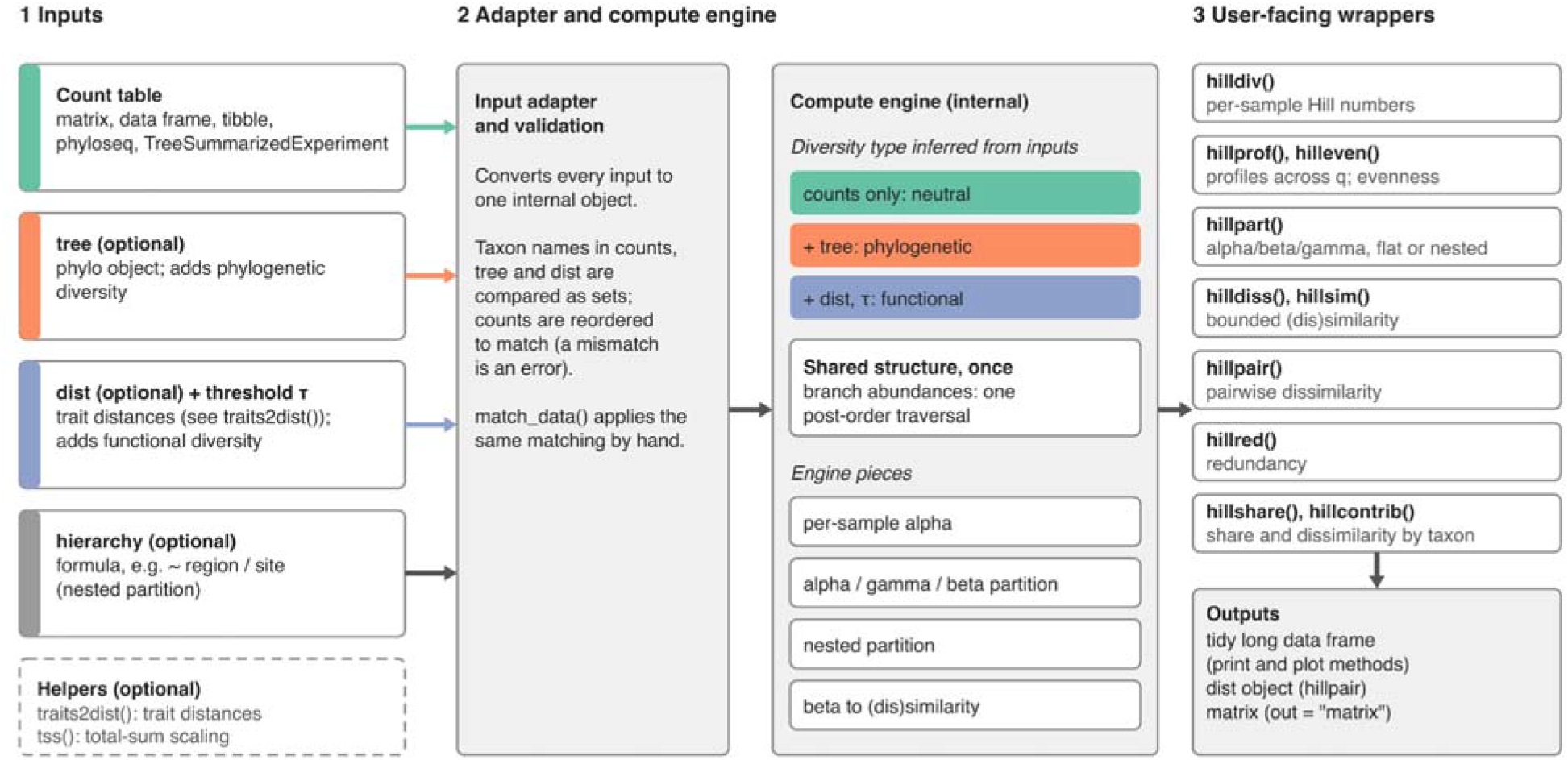
Structure of hilldiv3. All inputs (count tables as matrices, data frames, tibbles, phyloseq or TreeSummarizedExperiment objects, plus an optional phylogenetic tree, trait distance matrix with threshold τ, and hierarchy formula) pass through one adapter and validation layer into a shared compute engine. The engine infers the diversity type from the inputs (counts only: neutral; with a tree: phylogenetic; with a trait distance: functional), computes shared structure once, and provides per-sample alpha, alpha/gamma/beta partitioning, nested partitioning and the conversion of beta to (dis)similarity. Thin user-facing functions expose the engine and return tidy data frames, dist objects or matrices.

### Implementation

#### Diversity indices

A Hill number of order q expresses the diversity of a set of types (e.g., ASVs, MAGs, species) as an *effective number* of equally abundant, maximally distinct units (Hill, 1973; Jost, 2006). The order q sets the sensitivity to relative abundance: q = 0 ignores abundances and returns richness; q = 1 weights types by their frequency and equals the exponential of Shannon entropy; and q = 2 emphasises dominant types and equals the inverse Simpson index. Because every order is expressed in effective numbers of types, values are directly comparable across orders and obey the replication principle: doubling a system’s distinct content doubles its diversity (Hill, 1973; Jost, 2006).

The construction extends beyond the neutral case, in which all types are equally distinct. Phylogenetic Hill numbers replace types with the branch segments of a phylogeny, each weighted by its length and by the summed abundance of its descendants. This returns the effective number of equally abundant, equally distinct lineages at a chosen tree depth (Chao et al., 2010). Functional Hill numbers replace evolutionary distance with trait dissimilarity, returning the effective number of functionally distinct units at a distance threshold τ (Chiu et al., 2014).

hilldiv() computes Hill numbers per sample, by default at q = 0, 1 and 2, in whichever domain the inputs imply: counts alone give neutral diversity, counts plus a tree give phylogenetic diversity, and counts plus a trait distance matrix give functional diversity. The domain can also be asserted explicitly through type (auto-detected by default), which validates the inputs against the intended diversity type. For phylogenetic and functional diversity, the *reference* argument selects whether all samples are read at one common, mutually comparable tree depth or threshold (the default) or each at its own, and tau sets the functional distance threshold. Phylogenetic diversity is computed with a single post-order tree traversal (via ape (Paradis et al. 2004)) rather than repeated per-tip descendant look-ups, the principal source of its speed-up over earlier implementations.

#### Diversity partitioning

Hill numbers enable a clean and intuitive decomposition of diversity into within-sample (alpha), among-sample (beta) and total (gamma) components, as defined by Whitakker (Whittaker, 1960). Gamma is the Hill number of the pooled community, alpha is computed from the averaged basic sums of the samples (not by averaging their Hill numbers), and beta is obtained as gamma/alpha. So defined, beta is independent of alpha for all q, and ranges from 1 (all samples identical) to N (all samples wholly distinct). This value is read as the effective number of completely distinct communities, making it comparable across datasets that differ in richness (Jost, 2007). The decomposition carries over to the phylogenetic and functional domains using the same weighted-alpha construction, differing only in the per-feature measure and its normaliser (Chao et al., 2014; Chiu et al., 2014).

Real sampling designs, however, are rarely flat: samples are often nested within a hierarchy, such as replicates within sites, and sites within regions, and a single alpha/beta split collapses that structure into one number. The multiplicative decomposition of diversity generalises to such designs (Gaggiotti et al., 2018; Jost, 2007), provided that all levels share a common reference: rather than re-deriving weights at each level, every sample is given a single global weight (1/n under equal weighting, or user-supplied), each unit is weighted linearly by the sum of its sample weights as in a Routledge/Tsallis-type decomposition, and the beta at each scale is formed as the ratio of that scale’s Hill number to the next finer one. Under this construction, every beta is at least one for q ≥ 0, including in unbalanced designs, and the chain telescopes exactly (in balanced equal-weight designs each beta is also independent of alpha and bounded by the number of child units it aggregates): adjacent numerators and denominators cancel so the product of all per-level betas times the finest alpha returns the overall gamma. The same logic holds in the phylogenetic and functional domains, provided a single tree depth (or distance threshold τ) is fixed across all scales so the per-feature measure and its normaliser are shared throughout the hierarchy. This is a deliberate choice rather than a limitation: letting depth or τ vary by scale would break both the exact telescoping and the lower bound of one.

hillpart() returns the multiplicative partition for neutral, phylogenetic and functional diversity from the same engine. Supplying a *hierarchy* formula (e.g. ‘∼ region / site’) together with a metadata table switches it into nested, multi-scale mode: it builds the sample grouping implied by the formula and returns the finest-scale alpha together with one beta per hierarchical transition, plus the overall gamma. The three diversity types share this one construction, differing only in the per-feature measure and grand-total normaliser: neutral uses taxa (unit measure 1, normaliser n); phylogenetic uses branches weighted by length under one shared tree depth; and functional uses taxa weighted by their attribute contribution at a distance threshold τ. The nested result is returned as a dedicated ‘hill_hierarchy’ object with its own `plot` method. Samples can be weighted equally (the default), in proportion to their totals, or by a user-supplied vector; the phylogenetic depth can be set as a reference time on ultrametric trees, and a grid of τ values returns one hierarchy per threshold. Hierarchical output also feeds the (dis)similarity functions: hilldiss() and hillsim() return the overlap metrics for each hierarchical transition, using a data-dependent effective number of units that equals the child count in balanced equal-weight designs and keeps the metrics within [0, 1], and hillpair() computes pairwise dissimilarity between the units of a chosen level after weighted pooling of their samples.

#### (Dis)similarity metrics

The beta component produced by the decomposition above already encodes how distinct samples are from one another, but on a scale that runs from 1 to the number of samples N and therefore depends on how many samples were compared. This makes raw beta awkward as a (dis)similarity coefficient, which should lie in [0, 1] and be independent of N so that values are comparable across studies and usable by downstream statistics. The bridge from partitioning to dissimilarity is a set of simple monotonic transformations of beta that remove the dependence on N and yield similarity measures generalising familiar indices as special cases (Chiu et al., 2014; Jost, 2007). Two complementary families exist: Sørensen-type measures quantify (dis)similarity from the perspective of the individual sample, while Jaccard-type measures quantify it from the perspective of the pooled system. At particular orders, these reduce to classical coefficients such as the Jaccard and Sørensen indices (q = 0), the Horn index (q = 1) and the Morisita–Horn index (q = 2), and the same transformations apply to phylogenetic and functional beta, recovering measures such as unweighted UniFrac (Alberdi & Gilbert, 2019; Chiu et al., 2014). Dissimilarities are the one-complements of the corresponding similarities.

hilldiss() and hillsim() convert beta to bounded [0, 1] dissimilarity and similarity using the four overlap measures of Chiu et al. (Chiu et al. 2014), selected through the argument *metric*: the Jaccard-type (‘S’ and ‘U’) and Sørensen-type (‘C’ and ‘V’) measures. ‘S’ and ‘V’ are turnover measures, normalised to the pooled system (gamma) and to a single sample (alpha), respectively. ‘U’ and ‘C’ are the corresponding overlap complements: the proportion of non-shared units in the pooled system, and the effective average proportion of non-shared units in a single sample (‘C’ reduces to Morisita–Horn at q = 2). They operate in all three diversity domains, following the same type-inference rules as the rest of the package.

#### Pairwise (dis)similarities

Because these measures derive from the same beta diversity, they can be computed not only for a whole set of samples at once but for every pair. A pairwise dissimilarity matrix is the natural input to the multivariate methods that dominate community and microbiome analysis: ordination (PCoA, NMDS), hierarchical clustering, and distance-based hypothesis tests such as PERMANOVA. Grounding that matrix in Hill-number beta diversity means the ordination inherits the framework’s coherent properties: the order q tunes the abundance sensitivity explicitly, the same metric is available for neutral, phylogenetic and functional diversity, and the choice of coefficient (sample-vs system-perspective) is a deliberate, interpretable decision rather than an arbitrary one.

hillpair() returns the full matrix of pairwise dissimilarities as a dist object, ready for ordination, clustering and distance-based testing. It computes the type-specific shared structure (the pooled tree traversal or the trait distances) once and reuses it across all sample pairs, with optional parallelism over a future plan, keeping the all-pairs computation tractable for the large sample sets typical of high-throughput sequencing.

#### Shares and contributions

While beta diversity and (dis)similarity summarise how different communities are, two complementary quantities (share and contribution) address which taxa are responsible for the variation. A share allocates the Hill power sum of a single community, the quantity from which its Hill number is derived, among its taxa or among user-defined sets of taxa such as a phylum. In the neutral case, with relative abundance p_i_, the weight of taxon i is p_i_^q^ normalised by the sum over all present taxa, so that at q = 1 a share equals relative abundance. Phylogenetic weights are first assigned to branches (branch length multiplied by descendant abundance raised to q) and then divided among the descendant taxa in proportion to p_i_^q^, so that shared branches are apportioned rather than counted twice (Chao et al., 2010). Functional weights follow the same power sum with similarity declining linearly from one to zero at the threshold τ. Shares sum to one over a partition of the taxa within each non-empty sample and order, but they are fractions of the power sum, not of the effective number of units reported by hilldiv(). A contribution, in contrast, attributes a dissimilarity to taxa. For each taxon, or each phylogenetic branch, the kernel is the Jensen gap between its pooled abundance and its abundances across the compared samples, which is zero if the abundance is constant. The kernel fractions are multiplied by the chosen dissimilarity, so that taxon contributions add up exactly to the value returned by hillpair() or hilldiss(). For phylogenetic diversity, the gap of each branch is assigned to its descendant taxa in proportion to their neutral gaps at the same order, which is an explicit allocation rule rather than a unique decomposition of phylogenetic turnover.

hillshare() returns the share of every taxon, or of each named set supplied through sets, for every sample, order and diversity type implied by the inputs, using the same tree, dist, tau and type arguments as hilldiv(). Sets are lists of taxon names that may overlap, and the share of a set is the sum of the allocations of its members. hillcontrib() takes the same sets and returns, for every pair of samples (by = ‘pairwise’) or for all samples or named groups of samples together (by = ‘collective’, with groups), the contribution of each taxon or set to the dissimilarity, its fraction of that dissimilarity and its rank, for any of the four overlap metrics. Because taxon contributions are additive, aggregating them by sets such as phyla yields an exact budget at any taxonomic level, and sets that omit taxa sum to less than the total.

#### Complementary functions

Beyond a single diversity value, three further summaries help characterise communities. A diversity profile is the curve of Hill number values across a sweep of q; its shape (flat for even communities, steeply declining for uneven ones) conveys the entire abundance distribution at a glance and distinguishes communities that happen to coincide at one order (Alberdi & Gilbert, 2019; Jost, 2006). Evenness isolates the abundance-distribution component by expressing diversity relative to richness, and it ranges from 0 to 1. Redundancy captures how much phylogenetic or functional distinctiveness is lost relative to neutral diversity: when many co-occurring taxa are closely related or functionally similar, neutral diversity exceeds its phylogenetic or functional counterpart, and the gap measures the community’s redundancy.

hillprof() computes diversity profiles across a default sweep of q and hilleven() computes evenness, both in any of the three domains. hillred() estimates phylogenetic or functional redundancy by fitting the saturating relationship between neutral and phylogenetic/functional diversity across samples and summarising it as a single index per order. Two more helpers support all the aforementioned analyses: traits2dist() turns a trait table into the dissimilarity matrix required for functional diversity (Gower, Euclidean or Manhattan), and tss() performs total-sum scaling of a count table. The core verbs return tidy long-format output with ‘print’ and ‘plot’ methods, whereas hillshare() and hillcontrib() return plain long-format data frames.

### Comparative benchmarking

To test whether the generality of hilldiv3’s unified interface carries a computational cost, we benchmarked it against its predecessor hilldiv2 (2.0.5) and four widely used packages: hillR 0.5.2 (Li, 2018), entropart 1.6-16 (Marcon & Hérault, 2015), BAT 2.10.0 (Cardoso et al., 2015) and vegan 2.6-8 (Oksanen et al., 2013). The benchmark was based on a simulated dataset of 200 taxa and 50 samples, with ten timed iterations per package, carried out across four operations: (B1) alpha diversity at q = 0, 1, 2; (B2) diversity partitioning without a hierarchy; (B3) nested hierarchical partitioning; and (B4) pairwise beta-diversity matrices. Each operation was run in its neutral, phylogenetic and functional form wherever a package supported it, and wall-clock time and memory allocations were recorded with the bench package on a single machine (R 4.3.3, macOS 15.6, Apple silicon), using hilldiv3 3.2.0.

Of the twelve neutral/phylogenetic/functional operation combinations tested, hilldiv3 supported all twelve directly. hillR and hilldiv2 each supported nine, failing only on nested hierarchical partitioning (B3), which neither exposes. entropart returned a directly equivalent result for two combinations and a related-but-not-equivalent *partial* result for five more, as its phylogenetic and functional outputs rely on different normalisations. vegan and BAT, general-purpose diversity/ecology toolkits rather than Hill-number-specific implementations, only supported a single operation directly (neutral alpha), returning related-but-not-equivalent results elsewhere.

For the simplest and fastest case, i.e., neutral alpha diversity on a flat count table, alternative packages were faster than hilldiv3, owing to its fixed per-call overhead of input validation, alignment, and assembly of the standardised (S3) output, which dominates when the underlying computation is trivial (B1). However, as soon as heavier computation was needed due to the use of a phylogeny or trait distance, this ordering reversed. This is consistent with hilldiv3 evaluating tree- or distance-based quantities once and reusing them, rather than recomputing them across taxa, samples or pairs. The same pattern held for partitioning without a hierarchy (B2).

The advantage was largest, and most relevant to typical study designs, in nested partitioning and pairwise beta diversity. Nested hierarchical partitioning (B3) ran in 9–10 ms for all three diversity flavours, while the only package implementing a similar capability required approximately ten times longer. Pairwise beta-diversity matrices (B4) showed the sharpest contrast in both time and memory: hilldiv3 built the full neutral, phylogenetic and functional pairwise matrices in 54–70 ms using 45–98 MB, whereas hilldiv2 required 2.1–50.6 s (30–820× slower), peaking at over 20 GB of allocated memory for the phylogenetic matrix, and hillR required 0.8–16.2 s (15–230× slower) and up to 10.6 GB of allocated memory for the functional matrix.

**Figure 2.**
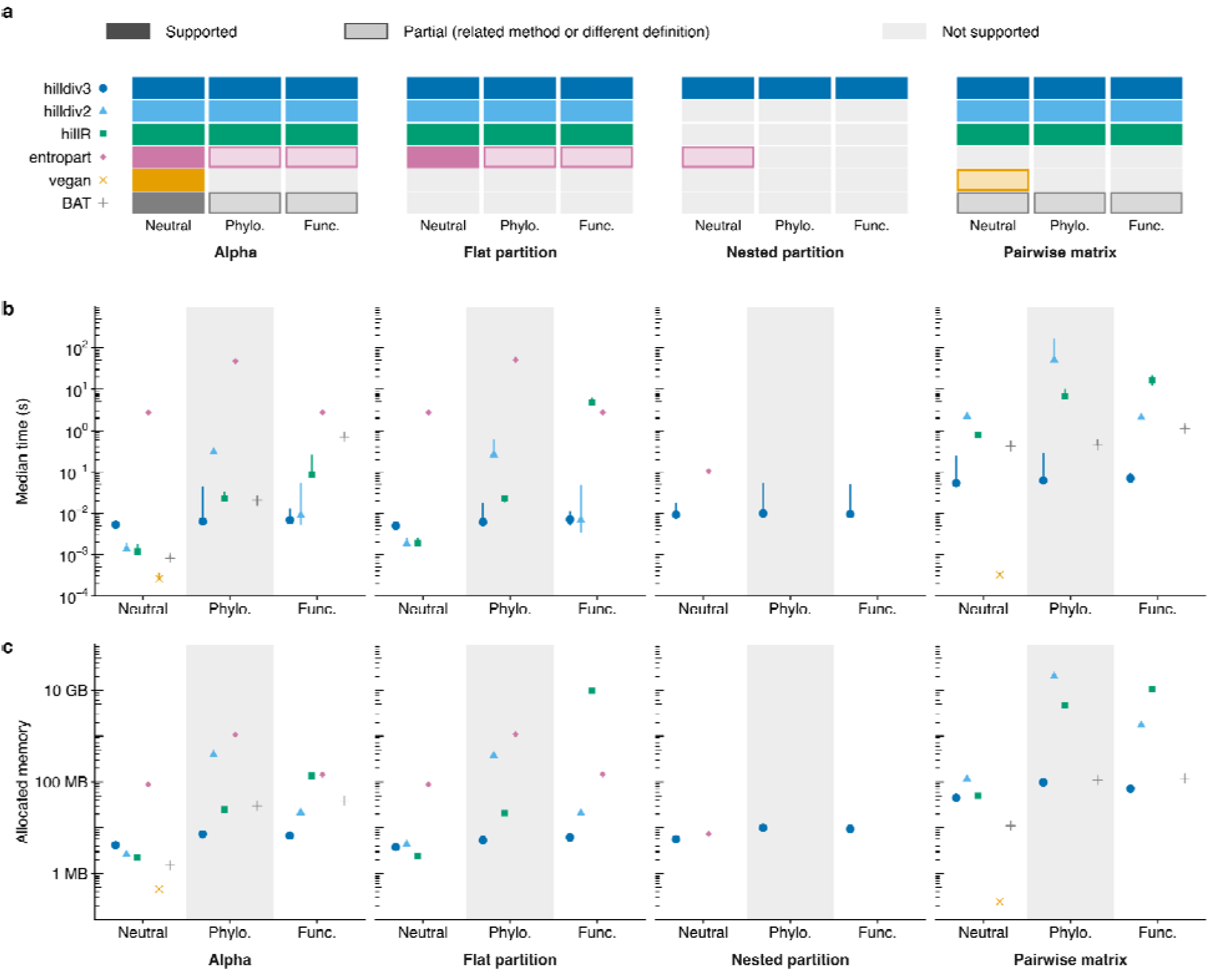
Performance of hilldiv3 compared with hilldiv2, hillR, entropart, vegan and BAT on simulated data. (200 taxa, 50 samples, 10 iterations). (a) Support for each operation (alpha diversity, flat partitioning, nested partitioning and pairwise beta matrices) in neutral, phylogenetic (Phylo.) and functional (Func.) form: supported directly, partially (related method or different definition) or not supported. The symbol next to each package name identifies it in panels b and c. (b) Median time (points) and minimum to maximum range (lines) on a log scale. (c) Memory allocated per evaluation on a log scale. Missing points are unsupported combinations.

### Use case: Danish environmental microbiomes

#### Data and analysis

To illustrate the framework on a realistic genome-resolved dataset, we analysed 360 samples from the Microflora Danica atlas of Danish environmental microbiomes (Singleton et al., 2026): 45 samples from each of eight habitat classes (agricultural field, natural grassland, natural forest, urban greenspace and bog/fen soils; freshwater and saltwater sediments; and urban wastewater).

The abundance table holds the published relative abundances of 5,518 species-level representative metagenome-assembled genomes (MAGs; 95% average nucleotide identity clusters), and the reference phylogeny was pruned to the same genomes. Functional traits represented 17 metabolic functions derived from KEGG ortholog annotations and converted to a Gower distance with traits2dist(). All analyses used hilldiv3 3.2.0 on this single abundance table, so moving between diversity types only required adding the tree or dist argument. Habitat is partly confounded with sampling year and the profiles are relative rather than count data, so the results are descriptive and no significance tests were applied.

#### Diversity across habitats

Within samples, median neutral diversity at q = 1 ranged from 47.8 effective MAGs in agricultural field soil to 78.0 in wastewater, which also had the highest phylogenetic (7.9) and functional (6.2) diversity (Table 1). Forest soil had the highest neutral diversity among the soils (74.5) but no matching increase in phylogenetic (4.8) or functional (5.3) diversity relative to field soil (4.9 and 5.3), so the additional MAGs there add neither lineages nor functions. Partitioning with hillpart() showed that the three diversity types turn over at very different rates. Neutral beta factors were 5.04 from samples to habitats and 3.33 from habitats to the full panel, against 1.46 and 1.56 for phylogenetic and only 1.06 and 1.03 for functional diversity. Comparing all 64,620 sample pairs with hillpair() gave the same ordering: median between-habitat C dissimilarity was 0.999 for neutral, 0.454 for phylogenetic and 0.07 for functional diversity. The habitats therefore consist of largely different MAGs, drawn from partly shared lineages, that carry largely overlapping functions.

**Table 1.** Hill-number diversity of the Microflora Danica panel at q = 1. Alpha is the range of habitat medians of per-sample diversity; its units (effective MAGs, effective lineages and effective functional types) differ among diversity types and are not directly comparable. β is the multiplicative beta factor from samples to habitats and from habitats to the full panel. C is the median pairwise C dissimilarity between samples from different habitats.

| **Diversity type** | **Alpha (range of habitat medians)** | **β samples to habitats** | **β habitats to panel** | **C between habitats** |
|---|---|---|---|---|
| Neutral | 47.8–78.0 | 5.04 | 3.33 | 0.999 |
| Phylogenetic | 4.4–7.9 | 1.46 | 1.56 | 0.454 |
| Functional | 4.3–6.2 | 1.06 | 1.03 | 0.07 |

#### Attributing turnover to taxa

Between the mean profiles of agricultural field and natural grassland soils, q = 1 C dissimilarity was 0.285 for neutral and 0.048 for phylogenetic diversity. Actinomycetota accounted for 37.9% and 35.3% of them, respectively (hillcontrib()), even though its share of the profile (hillshare()) was almost identical in the two habitats (34.9% and 34.7%). Beneath that stable total, Gaiellales declined from 12.5% to 3.0% of the profile while Streptosporangiales rose from 1.0% to 5.9%. To ask whether this swap changes what the community can do, we repeated the functional analysis with 161 finer-grained element-level traits, with τ set to the mean pairwise distance. Functional C between the same profiles was only 0.010, so most of the abundance turnover replaces MAGs with similar gene content. Yet the two orders, which make up 3.2% of the representatives, accounted for 19% of the neutral but 36% of the functional dissimilarity, and their contributions to element completeness were negatively correlated across elements (r = −0.78), indicating a partial offset rather than full functional equivalence. This excess fell to 18% (against 12% neutral) when restricted to higher-quality MAGs. From one abundance table, one phylogeny and one trait table, a single interface therefore separated extensive turnover in MAG identity from moderate turnover in lineages and slight turnover in function, and traced the contrast to specific clades.

**Figure 3.**
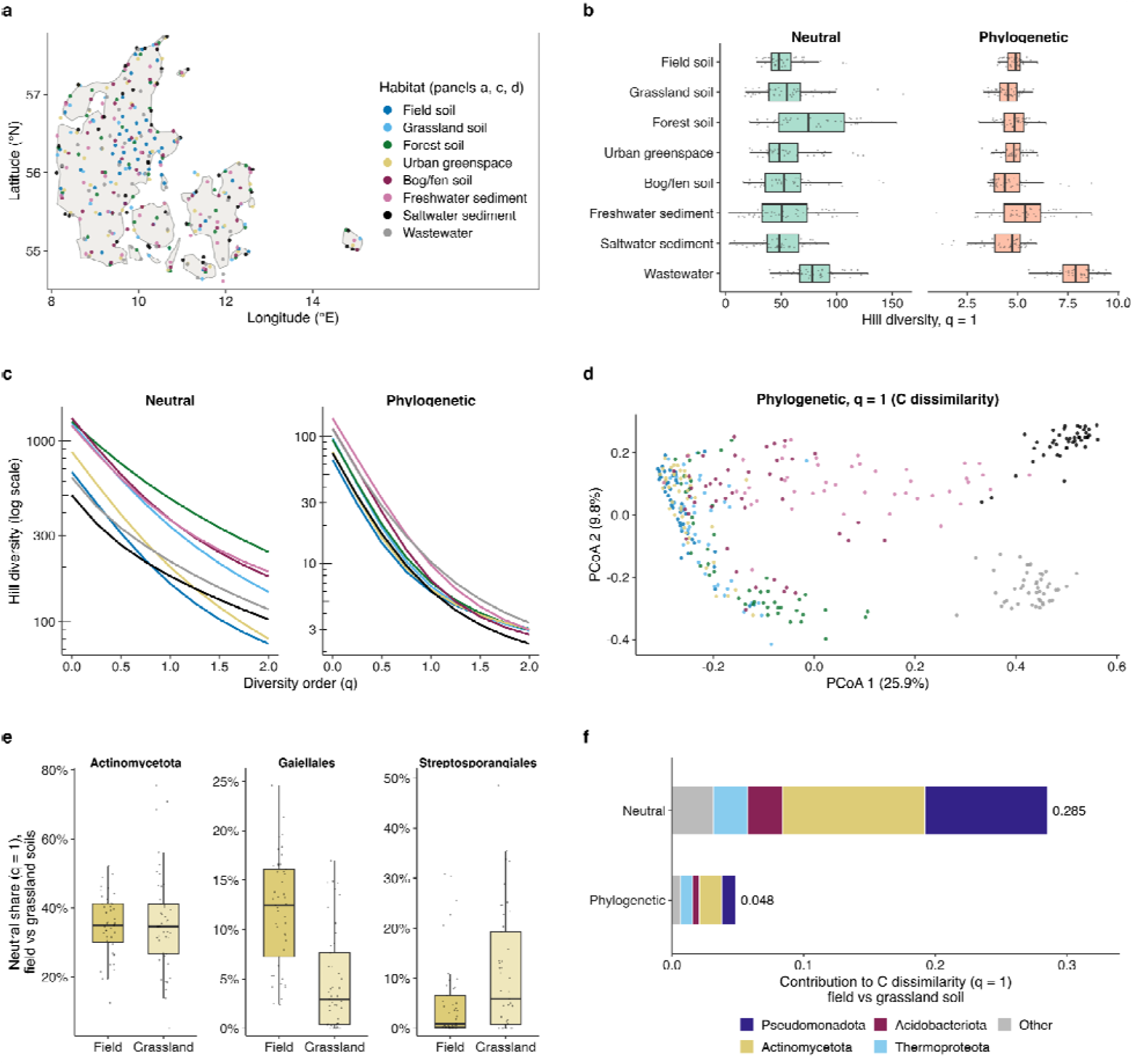
Hill-number diversity of the Microflora Danica panel analysed with hilldiv3. (a) Sampling locations, coloured by habitat. (b) Per-sample neutral and phylogenetic diversity at q = 1 by habitat (boxes: median and interquartile range; points: samples). (c) Neutral and phylogenetic diversity profiles across orders q = 0 to 2 for the mean abundance profile of each habitat. (d) Principal coordinates analysis of phylogenetic q = 1 C dissimilarity between samples, coloured by habitat; axes show the percentage of variance explained. (e) Neutral q = 1 shares of Actinomycetota and of its orders Gaiellales and Streptosporangiales in agricultural fields and natural grassland soils. (f) Partition of the q = 1 C dissimilarity between the mean field and grassland soil profiles across phyla, for neutral and phylogenetic diversity; totals are printed next to the bars.

## Discussion

‘hilldiv3’ provides a single route from a community table to neutral, phylogenetic and functional Hill numbers and to their partitioning, (dis)similarity, profiles, evenness, redundancy and taxon-level shares and contributions. Deriving every metric from one tested engine, and aligning inputs through one validation layer, makes the three diversity types and the full workflow mutually comparable. The Microflora Danica analysis showed what this buys: from one abundance table, one phylogeny and one trait table, the same interface separated large turnover in genome identity from moderate turnover in lineages and slight turnover in function, a contrast that is difficult to establish when each diversity type needs its own package and normalisation.

## Availability

‘hilldiv3’ is open source under the GPL-3 licence. Source code, issue tracker and documentation are available at https://github.com/alberdilab/hilldiv3 and https://alberdilab.github.io/hilldiv3/. It is available from CRAN (‘install.packages(“hilldiv3”)’), and the development version can be installed with ‘devtools::install_github(“alberdilab/hilldiv3”)’. Code and data for the Microflora Danica use case are available at https://github.com/alberdilab/hilldiv3-ms, with a snapshot archived in Zenodo with DOI 10.5281/zenodo.24001411.

## Acknowledgements

We acknowledge the Danish National Research Foundation for its grant DNR143 “Center for Evolutionary Hologenomics”.

## Use of artificial intelligence

Claude Code (Anthropic) and Codex (OpenAI) were used as coding assistants during development of the hilldiv3 R package, including drafting and refactoring code, writing unit tests and documentation, and reviewing code. Claude was also used to help draft and edit parts of this manuscript. All AI-generated or AI-assisted code and text were reviewed, tested and edited by the authors, who take full responsibility for the content of the paper and the software.

